# Pia-FLOW: Deciphering hemodynamic maps of the pial vascular connectome and its response to arterial occlusion

**DOI:** 10.1101/2024.02.02.577749

**Authors:** Chaim Glück, Quanyu Zhou, Jeanne Droux, Zhenyue Chen, Lukas Glandorf, Susanne Wegener, Daniel Razansky, Bruno Weber, Mohamad El Amki

## Abstract

The pial vasculature is the sole source of blood supply to the neocortex. The brain is contained within the skull, a vascularized bone marrow with a unique anatomical connection to the brain. Recent developments in tissue clearing have enabled unprecedented mapping of the entire pial and calvarial vasculature. However, what are the absolute flow rates values of those vascular networks? This information cannot accurately be retrieved with the commonly used bioimaging methods. Here, we introduce Pia-FLOW, a new approach based on large-scale fluorescence localization microscopy, to attain hemodynamic imaging of the whole murine pial and calvarial vasculature at frame rates up to 1000 Hz and spatial resolution reaching 5.4 µm. Using Pia-FLOW, we provide detailed maps of flow velocity, direction and vascular diameters which can serve as ground-truth data for further studies, advancing our understanding of brain fluid dynamics. Furthermore, Pia-FLOW revealed that the pial vascular network functions as one unit for robust allocation of blood after stroke.

## Introduction

Unlike other organs such as the liver or kidney, the brain is vascularized from outside where most large cerebral arteries lie horizontally above the cortical surface within the subarachnoid space (pia) (1). The pial vascular network is of utmost importance, as it is the only source of blood supply to the neocortex and a key contributor to cerebrovascular resistance (1, 2). The skull (calvarium), a vascularized bone marrow with highly interconnected vascular channels facilitating the trafficking of immune cells directly to the brain (3-5). Yet, a detailed depiction of fluid dynamics in pial as well as calvarial vessels remains remarkably incomplete, mainly due to the lack of quantitative hemodynamic imaging systems.

Past work provided detailed reconstructions of brain vasculature with impressive geometrical precision attainable (6-8). While highly informative, these reconstructions originate from *ex-vivo* imaging and thus do not provide functional information. Currently, most hemodynamic imaging techniques exhibit trade-offs in terms of imaging speed, quantitation, and invasiveness. While advances in mesoscale imaging techniques such as laser speckle contrast imaging (LSCI), optical coherence tomography (OCT), and magnetic resonance imaging (MRI) have enabled monitoring of cerebral blood flow, these techniques suffer from limited spatiotemporal resolution and/or lack of quantitative measurements (9-11). On the other hand, the current state-of-the-art two-photon fluorescence laser scanning microscopy (TPLSM) allows for quantitative blood flow measurements in pre-selected vessels utilizing line-scanning with maximal detectable velocities limited by the speed of scanning systems (12, 13). Recently developed technology now shifts TPLSM to kilo-Hertz frame rates which comes with some advantages for hemodynamic imaging, however limited to two dimensions (12, 13). TPLSM requires a cranial window preparation, which prohibits concurrent imaging calvarial and cortical vessels. Ultrafast ultrasound localization microscopy captures brain hemodynamics down to the microvascular level. However, imaging the pial vasculature is not possible (14, 15), because the projection of ultrasound echoes in one plane renders hemodynamic measurements valid only for vessels aligned with the axial direction hence the method is suitable only for coronal field of views (FOV).

Here, we present a large-scale imaging platform (Pia-FLOW) capable of structural and quantitative hemodynamic imaging of pial and calvarial vascular networks. With this method, we successfully captured the vascular architecture and blood flow dynamics at a spatial resolution of 5.4 μm (full width at half maximum, FWHM) and frame rates up to 1 kHz. Using Pia-FLOW, we provide maps of the pial and calvarial vascular networks across cortex-wide FOVs, which include quantitative information on vessel diameter, blood flow velocity and direction. We applied Pia-FLOW to quantify cerebral hemodynamic changes occurring in the pial vascular network after middle cerebral artery occlusion and identified diameter and flow changes in the arterial inflow, pial collaterals and venous outflow with an accuracy beyond conventional imaging techniques. Pia-FLOW offers new opportunities to investigate the principles of brain fluid dynamics.

## Results

### Pia-FLOW design and performance in terms of blood flow measurements

The current study presents both an experimental setup and dedicated data processing pipeline for dynamic imaging of brain pial and calvarial vascular networks. A simplified diagram of the Pia-FLOW system is shown in Fig. 1a. The system is based on a large-scale microscope equipped with a 473 nm continuous wave (CW) laser as excitation source and a high-speed sCMOS camera, for fluorescence detection of intravenously injected beads (FMOY, orange-yellow fluorescent microspheres, 50 μl of 2x10^8^ beads/ml) (Fig. 1a) (16). In brief, the laser beam was expanded with a lens pair consisting of a convex lens and an objective to provide widefield illumination. Back-scattered fluorescence was then collected with the same objective, focused with a tube lens and recorded by the camera. Pia-FLOW was performed at frame rates up to 1 kHz (~400-1000 Hz) to measure blood flow speeds even in vessels with high flow rates (until 20 mm/s).

**Fig. 1:**
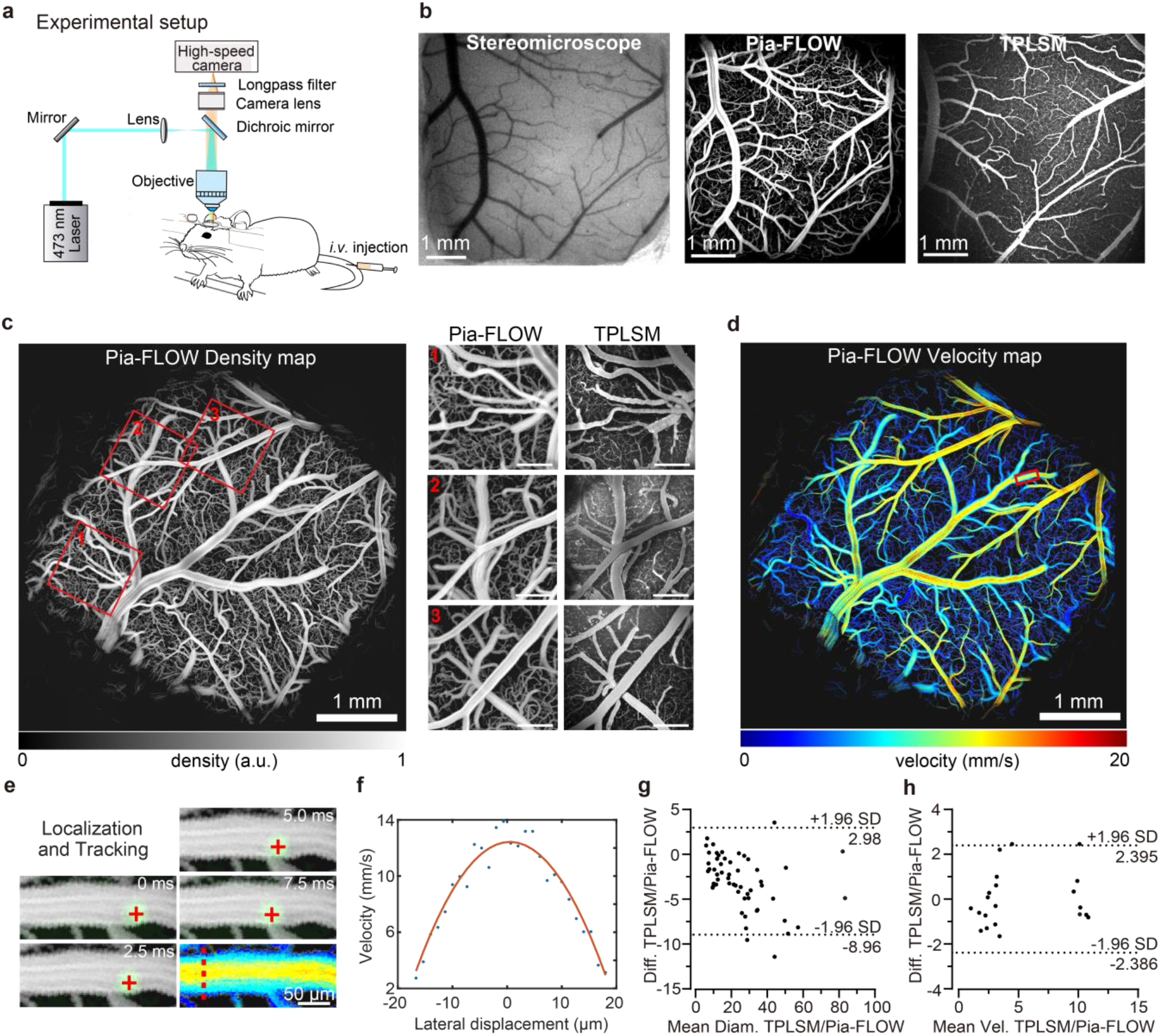
(a) Schematic of Pia-FLOW setup. **(b)** Stereomicroscopic, Pia-FLOW and TPLSM images of the same mouse brain vasculature. **(c)** Representative intensity map of the mouse brain with a cranial window recorded with Pia-FLOW post fluorescent beads injection. The red squares (1-3) indicate zoom-in views to compare the structural map acquired with Pia-FLOW to TPLMs (on the right). **(d)** Color-encoded flow velocity map. **(e)** Time-lapse images of a flowing fluorescence bead in the region of interest (ROI) indicated with a red square in (d). **(f)** Flow velocity along the red line labelled in (e) plotted as a function of the distance from the vessel center, fitted with a blunted parabolic function. (g-h) Bland Altmann analysis of Pia-FLOW and TPLSM for vascular diameters in µm (**g**) and blood flow velocity in mm/s (**h**).

Given that TPLSM is the most popular method for *in vivo* imaging, we imaged the same mouse brains with a 3×3 mm cranial window using both TPLSM and Pia-FLOW to compare the spatial resolution and accuracy of velocity estimation. First, we obtained a volumetric image stack of 200 µm using TPLSM covering all pial vessels to generate a maximum intensity projection (MIP) (Fig. 1b, c). Using Pia-FLOW, we generated high-resolution reconstructions of the pial vascular network with high-quality depictions of the vascular topology down to a 5.4 μm (full width half maximum; FWHM) resolution (Fig. 1b, c). In the intensity maps, fine details of vascular morphology were evident even in the smallest capillaries (Fig. 1b, c). We also obtained a similar signal intensity and angiography from TPLSM. Co-registration between Pia-FLOW and TPLSM revealed that both tools resolved the vascular architecture with nearly identical contrast and resolution for vessels with diameters larger than 10 µm. In contrast, we noted an advantage of TPLSM in regard to brain capillaries < 5 µm as only limited structural details were discernible in the Pia-FLOW configuration, potentially attributable to the spatial resolution of Pia-FLOW.

With Pia-FLOW, our goal was not only to achieve a robust *in vivo* mapping of the vascular architecture, but also to quantify flow rates and vascular diameters. Towards this goal, we utilized a particle tracking algorithm to extract the motion of fluorescent beads in the blood circulation. Typically, we performed continuous recording for 60-180 s, corresponding to 25,000-180,000 frames. Capitalizing on the precise temporal tracking of the sparsely distributed particles, we accurately extracted blood flow velocities and vessel diameters with a large dynamic range (from 0 to 20 mm/s and 5.4 μm to several hundred µm, respectively) (Fig. 1d, e). Representative time-lapse images of a single particle flowing in a selected vessel are depicted in Fig. 1e with a 2D velocity map reconstructed. The parabolic velocity profile across a large vessel is expected (Fig. 1f) and is in good agreement with the literature (13, 17). Since the number of particles that appear at each pixel was random, we averaged the flow velocity and direction values of different particles flowing across each pixel accordingly. Superimposing a sequence of images, we obtained high-resolution velocity maps of the entire pial vasculature (Fig. 1d, Supplementary Video 1).

To validate the accuracy of Pia-FLOW in quantifying hemodynamics, we selected vessels with different sizes inside the cranial window and performed TPLSM diameter and velocity line scans along the selected vascular segments. The Pia-FLOW estimates of blood flow speed and diameter at rest, are in reliable agreement with measurements made using TPLSM in the cortex. Bland-Altmann plots show a good agreement between the two methods over a wide measurement range (Fig. 1g-h). A significant advantage of Pia-FLOW is the ability to perform simultaneous flow measurements across different vessels. A large network of pial vessels can be scanned within 1-3 min. An *in vivo* example of this hemodynamic recording, with the resulting blood flow speed and directions is shown in Supplementary Videos 1 and 2. In contrast, imaging the same vessels with TPLSM took several hours. Overall, these results confirm that Pia-FLOW provides rapid simultaneous structural mapping and accurate hemodynamic quantifications with high spatio-temporal resolution.

### Transcranial hemodynamic imaging using Pia-FLOW

To apply Pia-FLOW for whole cortex imaging, it is critical to be able to perform recordings not only in mice with implanted cranial windows but also through the intact skull. Towards this goal, we further enlarged the FOV to 11×11 mm^2^ by using a different combination of objective and tube lens to image the entire brain transcranially. We reconstructed detailed structural maps of the entire pial microvascular network in both intact skull and cranial window implanted mice for comparative analysis (Fig. 2a-i). The excellent spatial resolution along the whole brain (Fig. 2g-i) enabled us to easily distinguish the three major arteries supplying the cerebrum (anterior, middle, and posterior cerebral arteries, ACA, MCA and PCA). Furthermore, we compared flow speeds from vessels imaged through a cranial window (Fig. 2a-d) to those imaged transcranially (Fig. 2e-h). Based on the vessel diameter and flow direction, vessels were divided into three groups: arteries/arterioles, veins/venules and capillaries. (Fig. 2i, j). We found that the transcranial velocities were in excellent agreement with the values measured through a cranial window (Fig. 2k, Suppl. Fig. 1). It is noteworthy that enlarging the FOV of Pia-FLOW to cover the entire pial network allowed for quantitative blood flow imaging in ~ 4000 vessel segments at the same time.

**Fig. 2:**
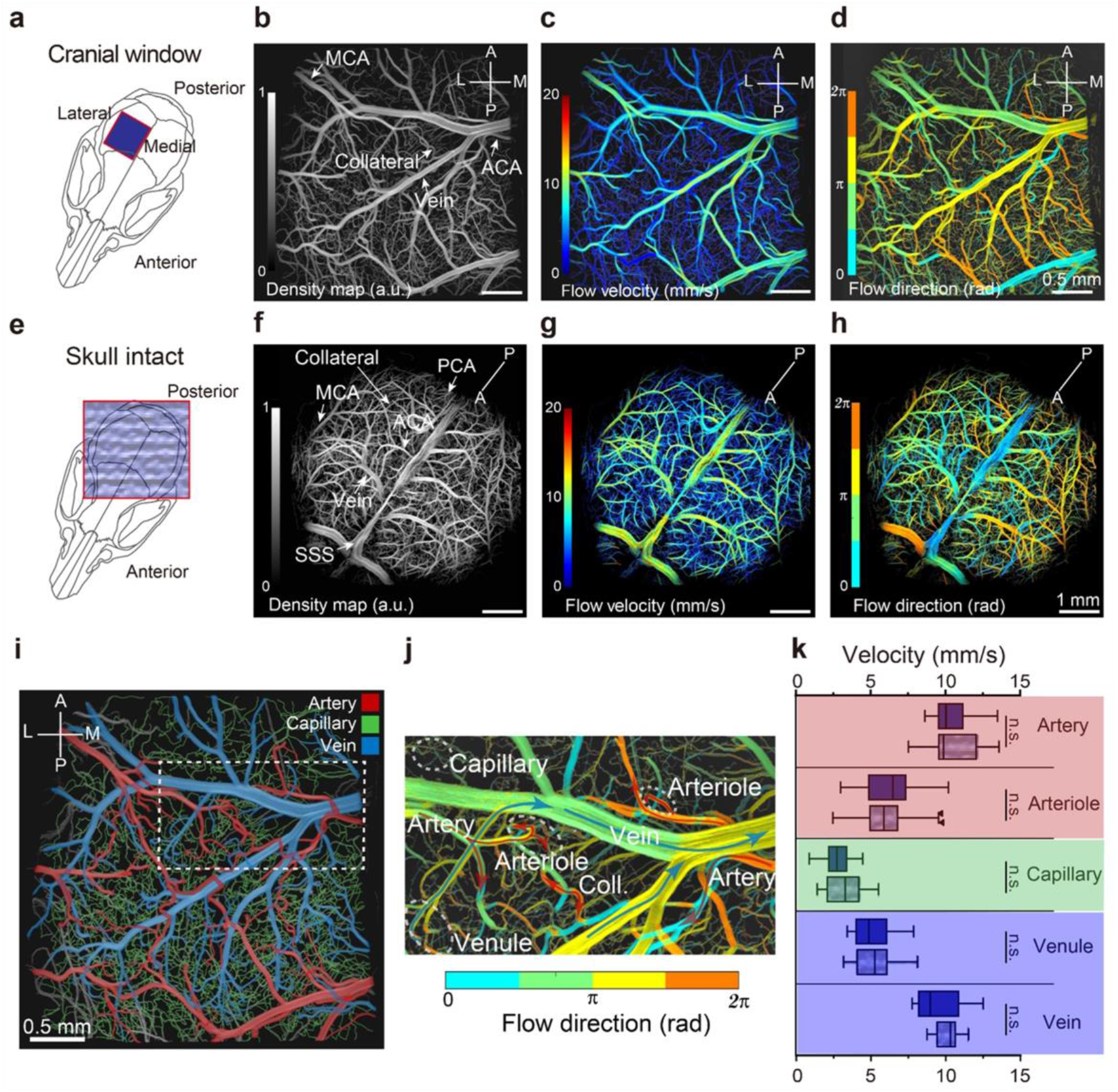
(a) Pia-FLOW measurement of a mouse with a cranial window implanted, showing **(b)** density map, **(c)** blood flow velocity and **(d)** directionality. **(e)** Pia-FLOW measurement performed transcranially, showing **(f)** density map, **(g)** blood flow velocity and **(h)** directionality. **(i)** Segmentation of arteries (red), veins (blue) and capillaries (green) shown in (b) based on flow direction in (d). (**j**) Indication of flow direction of a zoom-in region (indicated by dashed white square in (i). **(k)** Quantification of blood flow velocities in arteries, arterioles, capillaries, venules and veins comparing cranial window (dark blue) with transcranial (light blue) Pia-FLOW measurements. N = 3 mice for each group. Unpaired t-tests, n.s. = not significant.

In lower order arterial branches (100-200 µm), the measured blood flow speed was the highest and decreased progressively in higher order arteries and arterioles (50-100 µm) to reach the lowest rate in pial collaterals (terminal arterial branches) (Fig. 2c,g and k). Collaterals were easily observable as tortuous segments cross-connecting arterial branches with opposing flow directions: right to left (in orange) and left to right (in blue) (Fig. 2d, h). Fig. 2j shows the color-encoded flow direction map allowing the differentiation between veins, arteries and collaterals (Supplementary Video 2).

### Hemodynamic imaging of the calvarium using Pia-FLOW

Next, we evaluated the ability of our method to measure blood flow dynamics in the skull. The skull bone marrow is considered as the immune cell reservoir for the brain and calvarial vessels act as gates for the blood exchange between skull marrow and brain. To date, many fundamental questions regarding the vascular architecture and blood flow in the calvarium remain unresolved. The transcranial imaging capabilities of Pial-FLOW enabled us to conduct hemodynamic imaging in calvarial vessels, beyond the reach of conventional angiographic methods such as TPLSM (Fig. 3a-c). Using Pia-FLOW, we constructed the first velocity map of the murine calvarial vessels by the intravenous injection of Cy5.5 fluorescent dye followed by fluorescent beads. The time-to-peak (TTP) map generated from the dye perfusion process (Fig. 3d) served as a benchmark for segmenting calvarial vessels from pial vessels on the structural map reconstructed from Pia-FLOW (Fig. 3e, f). Thus, resolving fine details of calvarial vessels, we measured an average flow velocity of 4.7± 1.4 mm/s in the frontal plate, 5.4± 1.6 mm/s at the sagittal sinus interface; and 5.6± 1.6 mm/s at the parietal plate (Fig. 3g-i). Overall, our data demonstrate the capability of Pia-FLOW to perform simultaneous pial and calvarial hemodynamic imaging at single-vessel resolution.

**Fig. 3:**
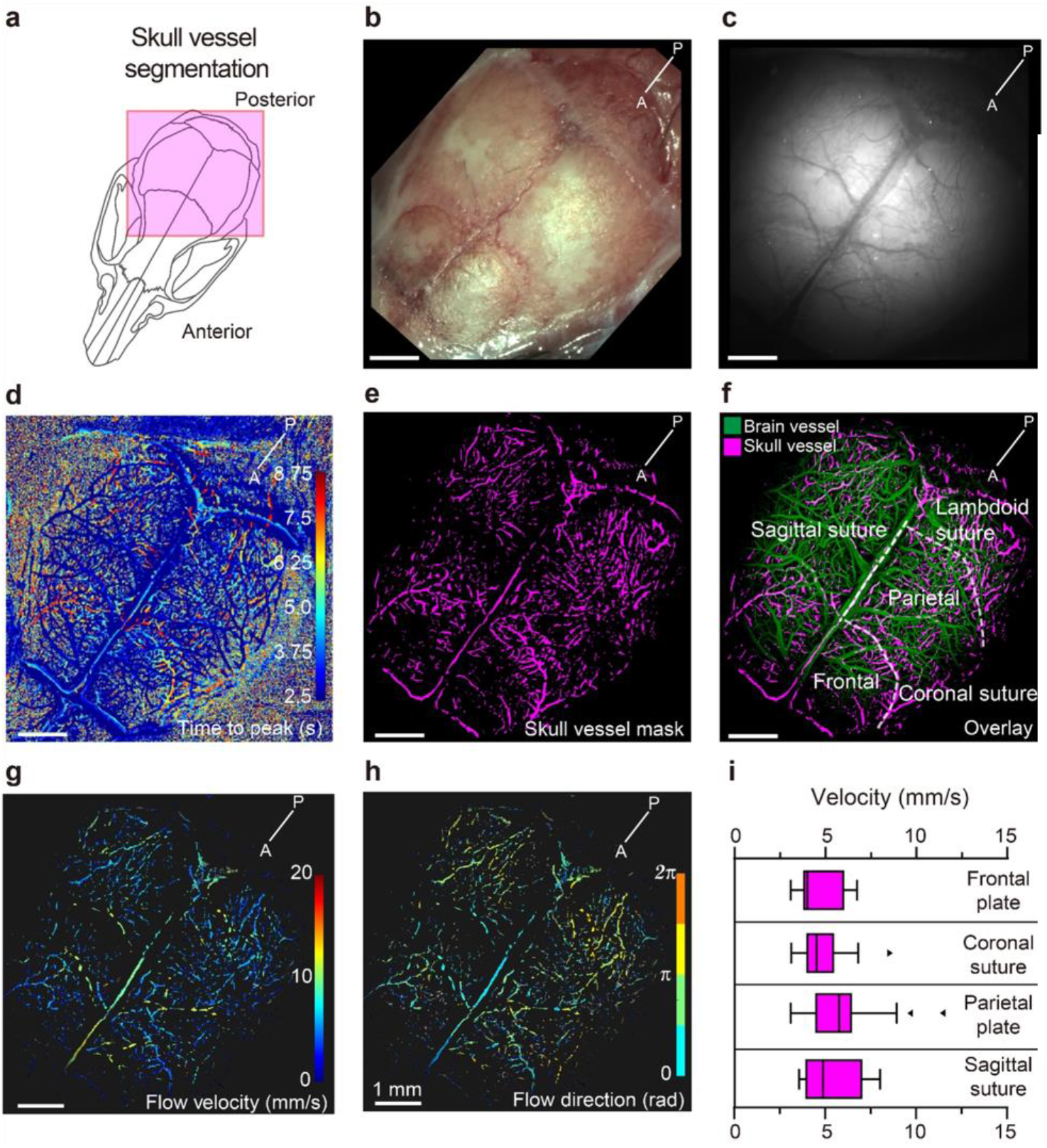
(a) Schematic of the skull vessel area imaged. **(b)** Exposed skull surface. **(c)** Skull illuminated with wide-field microscopy. **(d)** Time to peak (s) map of vessel perfusion upon dye injection. **(e)** Segmented skull vessel mask, based on time to peak map. **(f)** Overlay of skull vessel (magenta) onto pial vessels of the brain (green). Dashed white lines outline the skull plates and sutures. **(g)** Flow velocity and **(h)** flow direction map of skull vessels. **(i)** Quantification of the skull vessel blood flow velocity. N=4 mice.

### Measuring the response of the pial vasculature to MCA occlusion using Pia-FLOW

To further extend our technique to cerebrovascular research, we used Pia-FLOW to assess brain hemodynamics in the “thrombin” model of stroke in mice (Fig. 4a). Here, we compared Pia-FLOW with LSCI, which is another widefield imaging technology commonly used in stroke research. We collected datasets from three mice and each mouse was imaged with both modalities before and after stroke. In terms of spatial resolution, our technology demonstrated a clear advantage over LSCI by identifying some well-resolved vessel subtypes that remain undetected in the LSCI image (Fig. 4b, c). This difference became even more evident after stroke where the entire stroke area was blurred out, and LSCI failed to discern details on a single vessel level (Fig. 4c). To quantify flow changes in specific vessels, we used Pia-FLOW baseline images as reference and merged it with the corresponding LSCI image to select the same vessel segments for comparative analysis. We selected representative branches from arteries, arterioles, collaterals, venules and veins within the ischemic area. Pia-FLOW showed a reduction of blood flow in the highly interconnected arterial branches of the occluded MCA. The largest flow decreases (∼70%) were observed in high order arteries that were within a few branches downstream from the occluded segment (Fig. 4e). Lower-order (or more downstream) MCA branches (arterioles and penetrating arterioles) showed smaller flow rate reductions (Fig. 4e). Importantly, Pia-FLOW revealed that only collaterals showed significant increases in flow speed to redistribute blood supply to the affected arterial territory while the flow in ACA segments remained unchanged. These data are in line with previous TPLSM studies that discovered the engagement of collateral to reroute blood flow in stroke conditions (18, 19). In contrast to Pial-FLOW, LSCI was not sensitive enough to specifically detect the flow increase in collaterals but surprisingly showed flow reductions in both pial collaterals and ACA segments (Fig. 4e). Taken together, our results indicate that Pia-FLOW outperforms LSCI in terms of spatial resolution and accuracy in hemodynamic recordings.

**Fig. 4:**
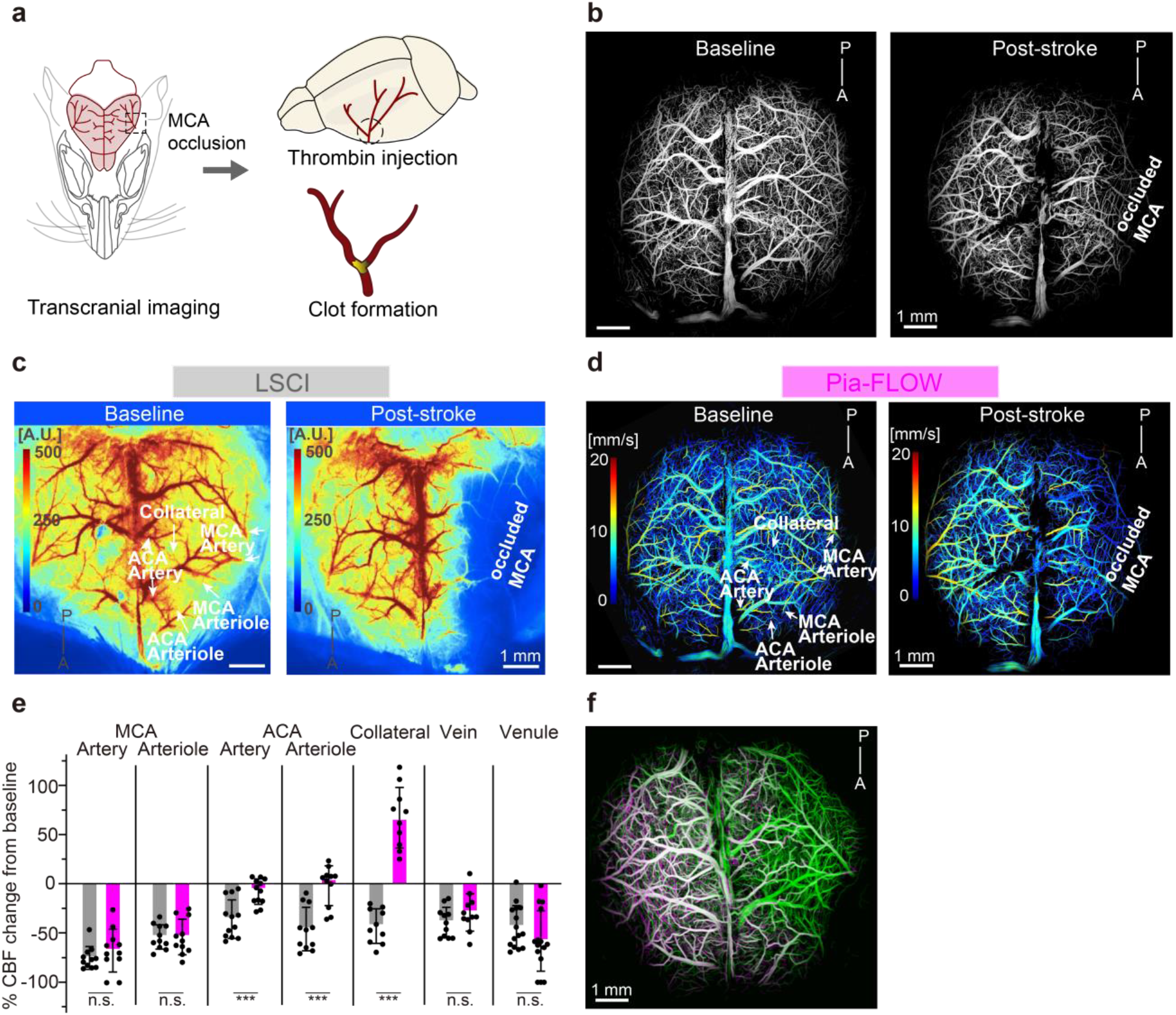
(a) Schematic of the stroke model. **(b)** Density images of a mouse at baseline and post-stroke. **(c)** Images of the same mouse brain acquired by LSCI and **(d)** Pia-FLOW at baseline and post-stroke showing perfusion or blood flow velocities. **(e)** Quantification of the change of blood flow in percentage of baseline. N=3, n.s. = not significant, *** P<0.001. **(f)** Overlaid image of a mouse brain acquired by Pia-FLOW showing vessel coverage at baseline (green) and post-stroke (magenta).

### The pial vasculature responds as a connectome after MCA occlusion

Next, we took advantage of Pia-Flow to investigate how each individual vessel in the pial network adapts in response to an arterial occlusion. Such experiments were previously difficult due to the need of simultaneous and quantitative measurements from the pial vasculature. For example, using TPLSM for this purpose takes several hours which renders capturing acute and transient effects impossible.

As pial collaterals are a natural bypass that can redistribute blood flow to ischemic regions, we primarily focused on these vascular segments. Previous work of us and others showed that after arterial occlusion collaterals dilate and change flow direction (18, 19). However, it is unknown whether all pial collaterals change flow direction after stroke and to what extent collaterals are capable of compensating blood flow in the affected region. Fig. 5a, b shows a specific vascular network before and after MCA occlusion. This network includes high and low order arterial segments from MCA, ACA and pial collaterals. First, we observed that under baseline conditions, collaterals show a preference in blood flow direction (some from MCA to ACA and others reversibly; Fig. 5 and Supplementary Fig. 2). The flow direction preference was defined by the higher flow gradient between MCA and ACA. Further, consistent with the idea that collaterals create a rescue road for blood supply we observed that after MCA occlusion collaterals divert blood from ACA to support the ischemic vascular network.

**Fig. 5:**
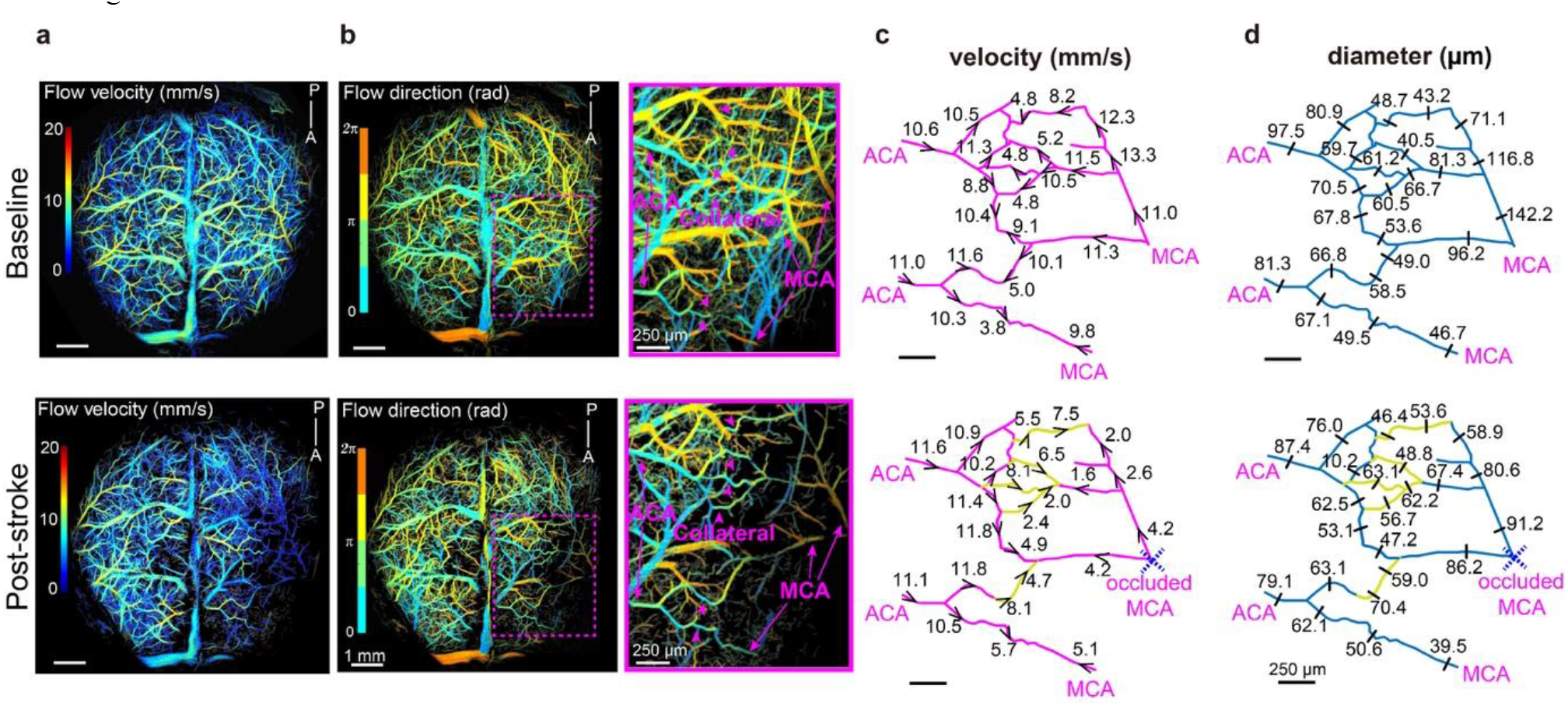
(a) Flow velocity map at baseline (top) and post-stroke (bottom). **(b)** Flow direction map. The zoom-in on the right (indicated by the dashed magenta square) highlights the position of collaterals (arrowheads). **(c, d)** Extracted arterial network. Values indicate (**c**) velocities and (**d**) diameters measured at indicated position with the arrowhead indicating the flow direction. Vessel segments with inverted flow direction post-stroke are indicated in yellow.

Interestingly, changes in blood flow direction occurred only in collaterals flowing originally from MCA to ACA. Altogether, our data show precisely how blood flow is rerouted from ACA via collaterals to MCA in order to compensate the flow reduction in ischemic stroke regions (Suppl. Fig. 2, Supplementary Videos 3 and 4).

## Discussion

Here, we presented Pia-FLOW, a mesoscale localization microscopy technique for structural and functional imaging of the pial and calvarial vasculature. We emphasize that Pia-FLOW concurrently achieves the following key benchmarks: (1) cortex-wide FOV, (2) transcranial imaging capability, (3) fast sampling rate (up to 1 kHz), (4) single-vessel resolution, and (5) quantitative measurements of flow velocity and vessel diameter.

We achieved a cortex-wide FOV not reachable with most of the available quantitative blood flow imaging tools. For example, conventional TPLSM is limited to sampling only a portion of the tissue accessible via cranial windows, unless a tiling strategy is used (20). To the best of our knowledge, no technology today comes close to being able to simultaneously track the entire pial vasculature at single-vessel resolution. Quantitative assessments of blood flow in the pial vasculature remained a major challenge impeding investigations of brain function and cerebrovascular diseases at large scale (21). Conventional widefield methods, such as laser speckle imaging, are limited with respect to their spatial resolution and only provide relative velocity measurements (22). Another key advantage of Pia-FLOW is its ability to perform quantitative and simultaneous blood flow measurements. TPLSM allows quantitative calculation of flow speed and diameters, however, this requires individual 1D line scans on each vessel making mapping the whole vasculature almost impossible. Although this problem could be overcome by using TPLSM Bessel beam, however, the maximum detectable velocity is around 3 mm/s constrained by the 2D frame rate of 99 Hz (23). Our widefield localization technique allows for the direct quantification of blood flow velocity (up to 20 mm/s) and diameters in hundreds of vessels synchronously. This can be particularly useful in cases where multiple vessels need to be measured synchronously in the whole brain vascular connectome. Outcomes from our work will supply the field with practical hemodynamic maps of the brain vasculature. Pia-Flow now allows systematic hemodynamic studies that will enhance our understanding of the pathophysiology of cerebrovascular disease.

Pia-FLOW is an easy and straightforward method that can be combined with existing widefield imaging systems used for fluorescence calcium imaging for neuronal activity. By incorporating these features, our microscope could be extended to synchronously measure blood flow and neuronal activity to study brain function in physiology and disease, such as neurovascular coupling and spreading depolarization.

Moreover, Pia-FLOW holds another key advantage because it allows imaging through the skull. This keeps the physiological environment of the brain intact. Previous studies showed that excision of calvarial bone to implement a cranial window may inadvertently remove parts of the dura (24) and also perturbates brain and vascular functions (25, 26). Although imaging through mechanically thinned skull bone reduces these problems (25), this may also cause trauma to trigeminal nerve terminals in the calvarium and inflammation of the dura by the axon reflex (27). Our study also demonstrates that Pia-FLOW can be used for noninvasive-transcranial imaging of the calvarial vasculature and provides topological, morphological, and hemodynamic information. This becomes increasingly important because of a growing interest in the skull marrow as a neuroimmunological niche (28). In fact, there is recent evidence that skull marrow is a myeloid cells reservoir for the central nervous system parenchyma (4). The skull vasculature provides signals for the maintenance and proliferation of hematopoietic stem and progenitor cells in the bone marrow (4). Previous work using optoacoustic ultrasound microscopy allowed imaging of the calvarial morphology but hemodynamic information was hard to achieve with this method (29). Thus, Pia-Flow is a promising tool that can be used to further study the vasculature in the skull marrow microenvironment to better understand how calvarial and pial vascular networks interact to regulate immune trafficking with the brain.

Using our method for proof-of-principle synchronous recording of the entire pial vasculature, we showed that after an arterial obstruction, the flow and diameter changes affect almost every pial vessel within the vascular network. More precisely, we took advantage of the ability of Pia-FLOW to perform cortex-wide simultaneous imaging and provided detailed insight into the spatial coupling of pial collaterals to the occluded and neighboring arterial branches. Our data showed that the whole brain vascular network react as a connectome where pial vessels redistribute blood to the ischemic territory via collaterals. Of note, such measurements would be incremental to understand recanalization/reperfusion and No-Reflow in stroke (30).

Finally, owing to the simple optical design, Pia-FLOW offers the potential to install an additional fluorescence channel for labeling other blood components such as leukocytes, red blood cells or plasma. We anticipate further important applications of Pia-FLOW, such as *in vivo* imaging of leukocyte trafficking in pial vessels, immune cells transition from the calvarium to the brain, and *in vivo* imaging of tumor metastasis.

Overall, Pia-FLOW achieves transcranial recordings of the pial and calvarial hemodynamics with unprecedented spatial and temporal resolution. Its ability to simultaneously measure blood flow in multiple vessels makes it a valuable tool to understand the control of blood flow in the brain. We expect Pia-Flow to become a widely used tool to investigate the brain and calvarial vascular biology and to find widespread applications in neuro-immunology and neurobiology.

## Material and Methods

### Ethics and Animals

Experiments were conformed to the guidelines of the Swiss Animal Protection Law, Veterinary Office, Canton of Zurich (Act of Animal Protection December 16, 2005 and Animal Protection Ordinance April 23, 2008, animal welfare assurance number ZH224/15 and ZH165/19). Experiments were performed on both male and female C57Bl6/J mice (Charles Rivers, no.028), 6 to 12 weeks of age, weighing between 20 and 30 g. The mice were housed under standard conditions including free access to water and food as well as an inverted 12-hour light/dark cycle.

### Anesthesia

For headpost and cranial window implantation, animals were injected intraperitoneally with a triple mixture of fentanyl (0.05 mg/kg body weight; Sintenyl, Sintetica), midazolam (5 mg/kg body weight; Dormicum, Roche) and medetomidine (0.5 mg/kg body weight; Domitor, Orion Pharmaceuticals). The face mask delivers 100% oxygen at a rate of 300 mL/min. For stroke induction, Pia-FLOW, TPLSM, and LSCI imaging, mice received the same triple anesthesia mixture intraperitoneally. During all procedures, the animals’ core temperature was maintained at 37 °C using a thermostatic blanket heating system (Harvard Instruments).

### Thrombin Stroke Model

To induce ischemic stroke, we used the thrombin model to induce stroke as previously described (31, 32). Briefly, a glass pipette was inserted into the lumen of the MCA and 1 μL of thrombin (1 UI; HCT-0020, Haematologic Technologies Inc) was injected to induce *in situ* clot formation (Fig. 3). To enhance the clot stability, the pipette was removed 10 min later.

### Scalp removal and cranial window preparation

After cranial window implantation, mice were allowed to recover for two weeks prior to two-photon imaging. For transcranial imaging, the skull of the mouse was kept intact while the scalp was removed to reduce light scattering. To minimize bleeding, hemostatic sponges (Gelfoam®, Pfizer Pharmaceutical) were used together with a topical application of adrenaline.

### Laser speckle contrast imaging

Cortical perfusion was monitored before and after ischemia using Laser speckle imaging monitor (FLPI, Moor Instruments, UK). The laser speckle images are generated with arbitrary units in a 32-color palette by the MoorFLPI software.

### Two-photon imaging

After cranial window implantation, mice were allowed to recover for two weeks prior to two-photon imaging. Imaging was performed using a custom-built two-photon laser scanning microscope (TPLSM)(33) with a tunable pulsed laser (Chameleon Discovery TPC, Coherent Inc., USA) equipped with a 25x (W-Plan-Apochromat 25x/1.0 NA, Olympus, Japan) water immersion objective. During measurements, the animals were head-fixed and kept under anesthesia as described above. To visualize the vasculature, Cascade blue Dextran (5% w/v, 10,000 kDa mw, 100 μl, D-1976, Life Technologies, USA) was injected intravenously 10 minutes before imaging and was excited at 820 nm. Emission was detected with GaAsP photomultiplier modules (Hamamatsu Photonics, Japan) fitted with a 475/64 nm band pass filter and separated by 506 nm dichroic mirror (BrightLine; Semrock, USA). The microscope was controlled by a customized version of ScanImage (r3.8.1; Janelia Research Campus(34). Line scans were processed with a custom-designed image processing tool box for MATLAB (Cellular and Hemodynamic Image Processing Suite (35); R2014b; MathWorks). Vessel diameters were determined at full-width-at-half-maximum (FWHM) from a Gaussian-fitted intensity profile drawn perpendicular to the vessel axis. Vessel flow was determined with the Radon algorithm.

### Pia-FLOW imaging

A 473 nm continuous wave (CW) laser (FPYL-473-1000-LED, Frankfurt Laser Company, Germany) beam was expanded with a convex lens (LA4306, Thorlabs, USA) and an objective (CLS-SL, EFL = 70 mm, Thorlabs, USA) to provide epi-illumination of the sample. 50 μl of fluorescent beads (FMOY, Cospheric, USA) in saline were injected via tail vein at a concentration of 2x10^8^ beads/ml. Backscattered fluorescence was then collected with the same objective, focused with a tube lens (AF micro-Nikkor 105 mm, Nikon, Japan), and detected by a high-speed camera (pco.dimax S1, PCO AG, Germany). A longpass filter (FGL515, Thorlabs, USA) was inserted between the tube lens and camera to filter out the reflected excitation light.

### Pia-FLOW image reconstruction and processing

To generate a high-resolution density, flow velocity and direction maps, the recorded widefield image stack was first denoised by subtracting a moving background calculated from the mean intensity of adjacent images to remove the static background stemming from the autofluorescence and dark noise of the camera. Sub-pixel localization was performed by extracting the centroid of detected bright spots detected using an adaptive threshold. Subsequently, the flowing fluorescent beads were tracked by identifying the same particle in different frames with simpletracker algorithm (simpletracker.m. GitHub https://github.com/tinevez/simpletracker, 2019), with no gap gilling and maximum linking distance corresponding to a velocity of 20 mm/s. Functional parameters, such as flow velocity and direction, were calculated with the given frame rate and relative displacement of the same particle in consecutive frames. A high-resolution intensity map was generated by superimposing the trajectories of beads. Similarly, flow velocity and direction maps were computed by averaging the velocities of all particles flowing through the same pixel. All the image reconstruction and processing were performed with custom code in MATLAB (MathWorks, MATLAB R2020b, USA).

### Quantification and statistical analysis

Parametric statistics were used only if the data in all groups in the comparison were normally distributed. Statistical analysis was performed using the GraphPad Prism (version 8.0; GraphPad Software La Jolla, CA, USA). All statistical tests and group size (n) are indicated in the figure legends. Results were expressed either as mean ± sd. (standard deviation). Significance (p < 0.05) between two groups was calculated using unpaired Student’s t-test or paired t-test for normally distributed data, or with the Mann–Whitney test for data with non-normal distribution.

## ACKNOWLEDGMENT

The authors acknowledge grant support from the Swiss heart foundation, the Swiss National Science Foundation (SNSF), and the UZH CRPP stroke.

## Supplemental Figures and Videos

**Supplementary Figure 1:** Velocities and diameters for example vessel ROIs recorded through (a) a cranial window and (b) transcranial.

**Supplementary Figure 2:** Extracted arterial networks showing velocities and diameters measured at indicated positions for baseline and post-stroke recordings. Cyan colored segments indicate flow reversal post-stroke. (a) mouse 2 and (b) mouse 3.

**Supplementary Video 1:** Short high-speed camera recording of fluorescent beads color encoded for velocity and overlayed on rendered vasculature image.

**Supplementary Video 2:** Flow direction map indicating the flow direction of tracked beads in a vein (ROI 1) and artery (ROI 2).

**Supplementary Video 3:** Short high-speed camera recording of fluorescent beads color encoded for velocity and overlayed on rendered vasculature image pre-stroke induction. (Related to Figure 5).

**Supplementary Video 4:** Short high-speed camera recording of fluorescent beads color encoded for velocity and overlayed on rendered vasculature image post-stroke induction. (Related to Figure 5).

