## Supplementary figures and images for "Pia-FLOW: Deciphering hemodynamic maps of the pial vascular connectome and its response to arterial occlusion"

### Supplemental Figure 1

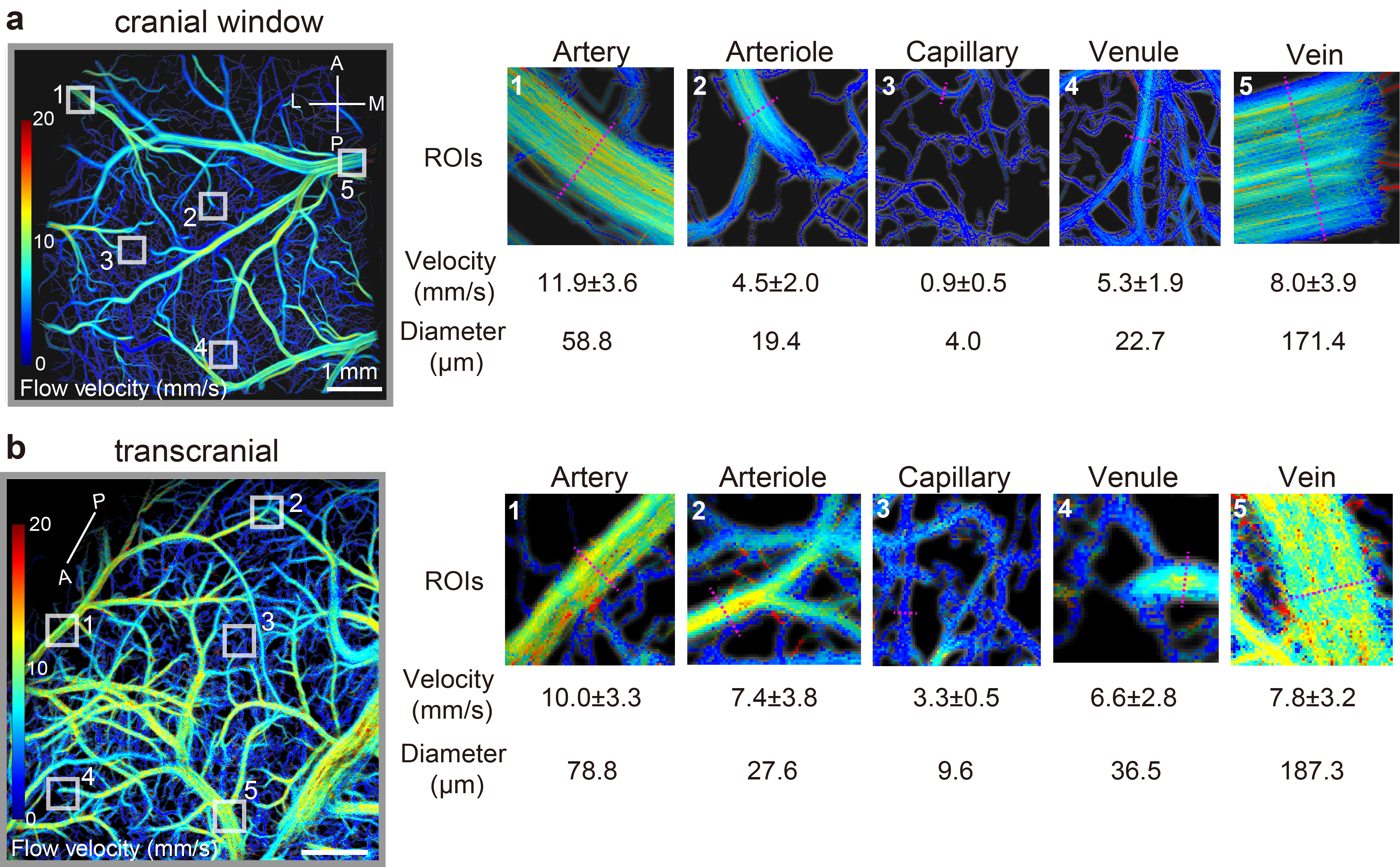

### Supplemental Figure 2

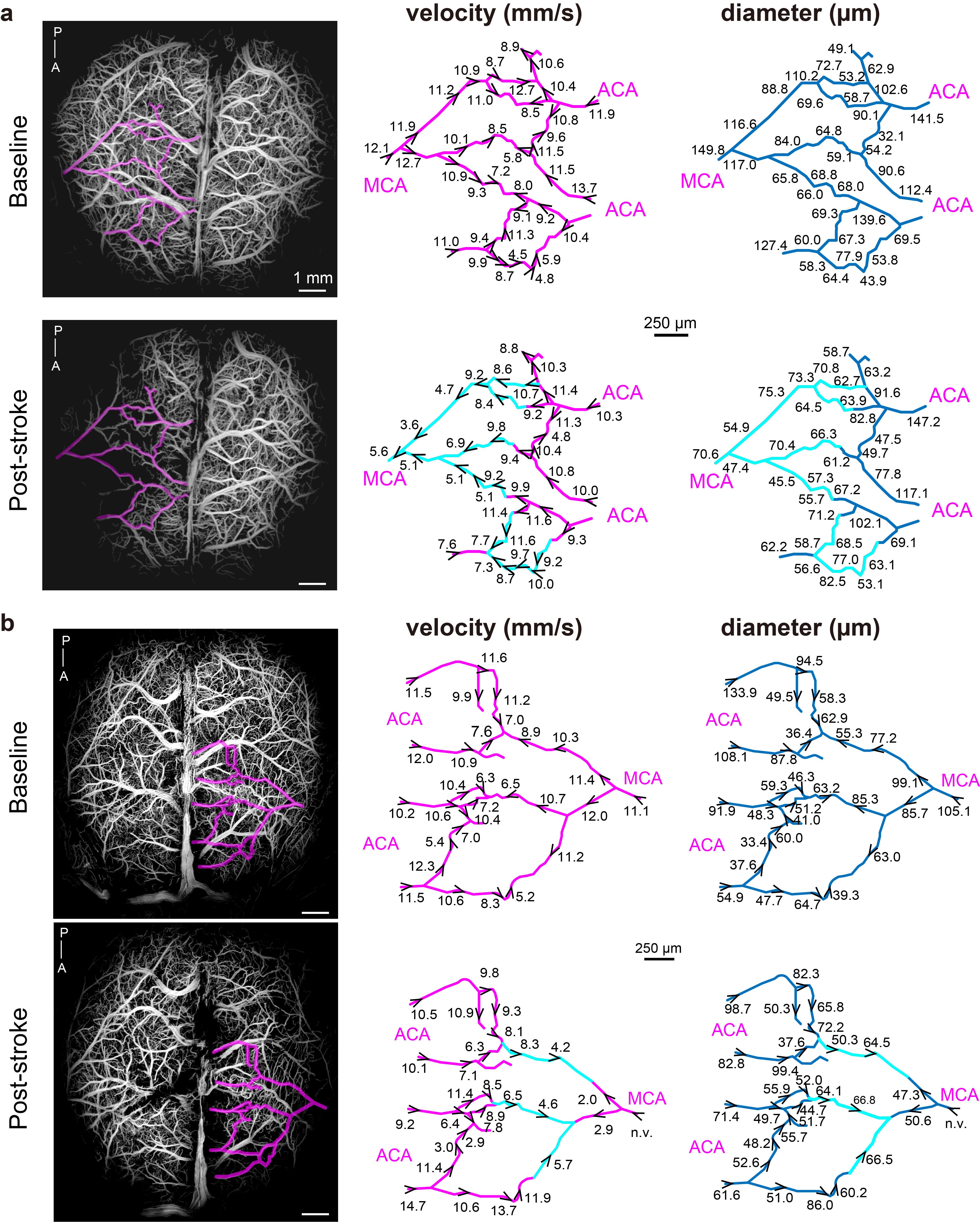
